# Long-acting naltrexone restores network connectivity in subjects with co-morbid cannabis and opioid use disorder

**DOI:** 10.1101/2025.09.18.676931

**Authors:** Lindsey M. Brier, Daniel D. Langleben, Corinde E. Wiers, Zhenhao Shi

## Abstract

Co-morbid substance use disorders (**SUDs**) are common but difficult to study due to the complex, interacting, and overlapping mechanisms through which they affect brain networks. Many datasets collected to investigate a specific SUD include participants with co-morbid SUDs. While most studies treat comorbid SUDs as covariates of no interest, these covariates also contain untapped information. This is particularly relevant as cannabis use disorder (**CanUD**) has become increasingly prevalent and co-morbid with other SUDs that have been more thoroughly studied. While treatments have been established for multiple SUDs, none have been approved for CanUD, although naltrexone (**NTX**) has been associated with reduced use. Here, we conducted a retrospective secondary analysis of functional magnetic resonance imaging (**fMRI**) data from individuals with primary opioid use disorder (**OUD**) with co-morbid CanUD, alcohol use disorder (**AUD**), or cocaine use disorder (**CocUD**), while controlling for opioid use. All participants underwent imaging prior to receiving a therapeutic dose of long-acting intramuscular NTX (Vivitrol^®^), an approved treatment for OUD and AUD but not for CocUD, and again two weeks post-administration. At baseline, OUD individuals with co-morbid CanUD, AUD, or CocUD exhibited distinct functional connectivity (**FC**) alterations compared to those with OUD-only. These differences were greater in younger participants and primarily involved the default mode network. Following NTX administration, FC differences between the co-morbid CanUD and OUD-only groups globally diminished. A similar FC response to NTX was observed in the parietal, subcortical, sensory, and cerebellar networks in the co-morbid AUD group. In contrast, little change in FC was observed in co-morbid CocUD. These findings, combined with prior evidence that NTX reduces cannabis use by dampening the experience of reward, suggest NTX may hold promise as a treatment for CanUD.

## 1. Introduction

Co-occurring and co-morbid substance use disorders (**SUDs**) are becoming the norm rather than the exception among individuals who misuse alcohol, cannabis, opioids, or cocaine, as well as those with a psychiatric disorder [1]. The etiology of these co-morbidities is likely multifaceted, involving impure street manufacturing processes, social environments and norms surrounding use, self-medication of symptoms or side effects from one substance with another, and attempts to maximize the psychoactive effects of any one substance [2]. A well-known example is the concurrent use of alcohol and cocaine, which produces cocaethylene, a psychoactive metabolite that, like cocaine, inhibits the re-uptake of dopamine but has a longer half-life, resulting in prolonged and intensified intoxication. Some users also combine alcohol (a depressant) with cocaine (a stimulant) to mitigate the undesirable effects of either substance when taken alone [3]. However, for other substance combinations, the perceived benefits to users are less clearly understood.

Among patients with opioid use disorder (**OUD**), approximately 28% have co-morbid cannabis use disorder (**CanUD**) [4]. The reasons for this co-occurrence remain unclear, though contributing factors include cannabis’ potential to reduce the severity of opioid withdrawal symptoms and the parallel rise of cannabis legalization and the opioid epidemic in the United States [5]. The phenomenology of OUD has evolved significantly since the 20^th^ century era of injectable heroin abuse as heroin has largely been supplanted by illicitly manufactured fentanyl, a synthetic opioid many times more potent than heroin and the leading cause of overdose deaths in the past decade [6]. Medication-assisted treatments (**MAT**) such as methadone and buprenorphine are effective in aiding detoxification and maintaining sobriety in OUD and have been utilized more frequently and for longer in response to spreading opioid use [7, 8].

Simultaneously, cannabis use in the United States has steadily increased. As of 2022, 25% of individuals aged 12 or older reported cannabis use, with 19 million meeting DSM-5 criteria for CanUD [5]. The legal cannabis industry is now valued at nearly $40 billion, with continued growth projected. This trend coincides with expanding state-level legalization and a declining perception of risk among the public [9]. In response, dispensaries are producing cannabinoid products with higher purity and potency, the most popular containing the dually psychoactive and addictive delta-9-tetrahydrocannabinol (**THC**). Subjectively, reported benefits of THC usage include feelings of relaxation, pleasure, and dissociation in the short term [5]. Objectively, research has indicated an increased incidence in cognitive and mood disorders, anxiety, and psychosis in the long term [10].

At the molecular level, prior works have begun to explore the neurobiological basis of OUD and CanUD comorbidity. Both opioid and cannabinoid (**CB**) receptors are G-protein coupled and co-localize on pre-synaptic terminals in limbic, mesencephalon, brain stem, and spinal cord regions [11]. The exact mechanism by which opioid and CB receptors interact is unknown, however some studies suggest that mu-opioid and CB1 receptors function together, as either heterodimers or allosteric modulators would [12]. Elsewhere in clinical studies, patients receiving both THC and opioid analgesics for treatment of chronic pain reported a 27% decrease in pain compared to opioids alone without any change in blood opioid concentration, concluding that THC augments the experience of opioids by some other mechanism [13].

As mentioned above, multiple FDA-approved treatments are available for OUD. Within the category of opioid substitution treatments are the MATs mentioned earlier, full opioid agonist methadone and partial opioid agonist buprenorphine. Currently, these medications are popular as they are initially used for detoxification from illicit opioid use and many patients elect to continue them as maintenance therapy [7, 8]. Although the recommended duration of MAT is unknown, it is measured in years, with early discontinuation highly predictive of relapse [14, 15]. The need for long term treatment and high risk of nonadherence led to the development of long-acting injectable preparations. Buprenorphine and opioid antagonist naltrexone (**NTX**) are both medications that come in longer acting injectable forms that can help mitigate this burden by requiring a once a month shot for MAT [16].

NTX is also an efficacious treatment for alcohol use disorder (**AUD**). Broadly, NTX inhibits B-endorphin activity by blocking opioid receptors in the ventral tegmental area and nucleus accumbens which leads to a decrease in dopamine release translating to reduced reward associated with substance use and therefore lower levels of use [17]. While NTX has been clinically proven to decrease levels of alcohol and opioid use, data have not supported NTX in the treatment of cocaine use disorder (**CocUD**) [18]. This may be due to cocaine producing a euphoric effect through dopamine re-uptake blockade as opposed to opioid receptor modulation [19]. Although NTX is known to reduce craving and reward of various substances [20], its benefit for CanUD is uncertain. However, a clinical study supplying long-acting NTX to cannabis users noticed a decrease in number of days of use and overall perceived reward from using within 2 weeks of being on NTX. This finding was maintained during the post study follow up and postulates that NTX could have some utility in treating CanUD or helping cannabis users cut back on use in the same manner it is effective in OUD and AUD, however further investigation is needed [21].

Functional magnetic resonance imaging (**fMRI**) is a non-invasive modality that has the unique ability to explore neuroscientific phenomena at both the molecular and systems level [22, 23]. In addiction research, fMRI has been used to describe a degradation in functional network segregation in adults of all ages with OUD and AUD, that scales with addiction severity and is similar to what would be expected in age-related cognitive decline [24, 25]. In CanUD, most neuroimaging studies focus on adolescent subjects. In this population, there is evidence for THC-induced altered hippocampal activity during memory encoding [26]. Also in adolescents, functional connectivity (**FC**) was noted to increase in dorsal medial prefrontal cortex and decrease in dorsal visual stream networks following acute THC consumption in chronic cannabis users [27], though no similar analysis has been done in adults. Functional readouts such as these are frequently used as biomarkers that track severity of illness as well as response to treatment and have been rigorously validated in disease processes such as Alzheimer’s disease [28], stroke [29], and multiple sclerosis [30].

Here, we analyzed resting state (**rs**) brain fMRI data that was collected in OUD patients before and after receiving NTX. Each participant had accompanying data on other substance use as well as co-morbid SUDs, allowing us to separate the data into OUD with co-morbid CanUD (**CanUD+OUD**), OUD with co-morbid AUD (**AUD+OUD**), and OUD with co-morbid CocUD (**CocUD+OUD**) compared against OUD-only (to control for opioid use). We used the Power atlas to define 264 regions of interest (**ROI**) and performed Pearson correlation analysis on the average fMRI time-trace within each ROI to calculate functional connectivity (**FC**) of each ROI permutation. Using this methodology, we commented on the functional changes introduced by co-morbid SUDs as opposed to OUD alone and monitored each combination of SUDs response to NTX. We hypothesized that NTX would mitigate network changes due to co-morbid AUD and CanUD, and this effect would be reduced in the presence of CocUD.

## 2. Methods

### a. Participants

Two datasets were used in the following analysis (imaging parameters for each dataset described below). For both datasets, treatment-seeking participants with OUD (heroin or pill opioids) were recruited in the greater Philadelphia area between 2012-2014 to receive up to three monthly extended-release NTX injections, as previously described [25, 31]. Both datasets were collected following protocols approved by the university’s Institutional Review Board, and all subject’s signed voluntary consent to treatment and imaging sessions.

A total of 69 subjects participated in the baseline imaging session (**pre-NTX**). Data quality screening removed subjects due to motion artifact (N=4; see criteria below), incomplete data (N=3), or FC matrices that were statistical outliers (N=2). A total of 60 subjects participated in the second imaging session (**on-NTX**). Data quality screening removed subjects due to motion artifact (N=1), incomplete data (N=3), or FC matrices that were statistical outliers (N=1). As the purpose of this analysis was to differentiate FC changes with respect to co-morbid SUDs on top of OUD, subjects with years of opioid use that were statistical outliers were excluded from the following analysis (N=5, SFigure 1A) at both time-points.

The same experimental timeline was used for both datasets. Initially, subjects underwent outpatient detoxification from illicit opioid use prior to the first MRI scan (pre-NTX). Following the baseline scan, subjects received the first intramuscular TX injection (380 mg) within the first week (N=44), within the next 2-6 weeks (N=4), or dropped out of the study (N=7). The second MRI scan was an average of 11.8 days (standard deviation 4 days) from the first NTX injection (**on-NTX**).

Following data quality exclusion as described above, a total of 55 adult human subjects were imaged and analyzed pre-NTX, with an average age of 27.7 years and standard deviation of 7.4 years and 63.7% male. A total of 50 adult human subjects were imaged and analyzed on-NTX, with an average age of 27.4 years and standard deviation of 7.4 years and 66% male.

For the purposes of this study, subjects were split into groups of either OUD-only, CanUD+OUD, AUD+OUD, or CocUD+OUD (SFigure 1B) at each time-point. Subjects who met criteria for more than two SUDs (e.g., OUD with co-morbid AUD and CanUD) were not used in the following analysis. The number of subjects included within each comparison is listed above the group-wise averaged FC matrix in each figure.

### b. Drug use severity

The Addiction Severity Index-5^th^ edition (**ASI**) [32] cumulative score served as a marker for drug use severity in each subject. Components of this metric represent factors such as medical status, employment and support, drug use, alcohol use, legal status, family/social status, and psychiatric status. A higher cumulative ASI score indicates a more severe recent drug use history.

### c. Image acquisition and pre-processing

The imaging data were collected on a Siemens Trio 3T scanner (Siemens AG, Erlangen, Germany). The first rs-fMRI dataset (previously described [25]) was collected using a whole-brain, single-shot gradient-echo echo-planar sequence with repetition time (TR)/echo time (TE) = 2000/30 ms, field of view (FOV) = 220×220 mm^2^, matrix = 64×64, slice thickness/gap = 4.5/0 mm, 32 slices, with effective voxel resolution of 2. 4×3.4×4.5 mm^3^, flip angle (FA) = 90°. The second rs-fMRI dataset was collected with the same settings except for TR/TE = 3000/32 ms, FOV = 192×192 mm^2^, matrix = 64×64, slice thickness/gap = 3/0 mm, 46 slices, with effective voxel resolution of 3×3×3 mm^3^, FA=90°. For structural imaging (previously described [33]), the magnetization-prepared rapid acquisition gradient echo sequence acquired high-resolution T1-weighted whole-brain images with TR/TE = 1510/3.71 ms, FOV = 256×192 mm^2^, matrix = 256×192, slice thickness/gap = 1/0 mm, 160 slices, with effective voxel resolution of 1×1×1 mm^3^, FA = 9°.

The rs-fMRI data were preprocessed in MATLAB using a pipeline adapted from Ciric and colleagues (2018) [34]. This consisted of removing the first five scans, estimation of the 24 motion parameters (including the six raw motion parameters, six framewise displacement (**FD**) parameters, the square of the raw motion parameters, and the square of the FD parameters), identification of FD timepoints (> 0.5 mm), removal of subjects with absolute maximum displacement from the first frame > 3 mm, identification of slice time correction, motion correction, coregistration and segmentation of the structural images, skull stripping, computation of DVARS and identification of DVARS outliers using the procedure described in Afyouni & Nichols (2018) [35], despiking using AFNI’s 3dDespike, removal of polynomial trends (order = 3), extraction of nuisance signals from the voxels located within the top 10% of the deepest tissue of the white matter and the cerebrospinal fluid, interpolation of FD and DVARS outlier timepoints using Lomb-Scargle periodogram, bandpass filtering at 0.01–0.1 Hz of the images and covariates (i.e., 24 motion parameters and two nuisance signals), regressing out the filtered covariates, spatial smoothing using a Gaussian kernel with a full width at half maximum of 8 mm, and spatial normalization to the Montreal Neurological Institute space.

### d. Functional connectivity analysis

The Power-264 atlas [36] was used to define a set of 264 5-mm-radius evenly distributed spherical ROIs and the average time trace within these ROIs was extracted. These ROIs were distributed amongst 13 networks: “association” networks (default mode, fronto-parietal, medial parietal, ventral attention, dorsal attention, cingulo-opercular, and salience); “sensorimotor” networks (hand sensory motor, mouth sensory motor, visual, and auditory); and “other” networks (subcortical and cerebellar). A Fisher-Z transformed Pearson’s correlation coefficient served as a measure of FC among the ROI permutations.

### e. Statistical analyses

Statistical comparison of FC matrices was performed at each time-point between the two groups described in each figure (either OUD-only vs CanUD+OUD, OUD-only vs AUD+OUD, or OUD-only vs CocUD+OUD) via pair-wise two-sample t-test. To correct the statistical matrix for multiple comparisons, we the statistical matrix gets iteratively thresholded (*h*) from zero to the maximum statistical value (thresholding set implemented a threshold-free network-based statistical (**TFNBS**) method, as described elsewhere [37]. Briefly, at 101 steps, *dh*). At each threshold, connected components are identified and each matrix index is replaced with the size of the component that index sits in (*e*(*h*)). Then the values within each index are summed across threshold steps to create the *TFNBS* matrix, whereas:

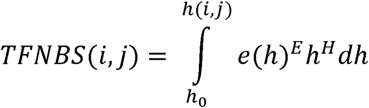

Where *i* and *j* index the statistical matrix, *e*(*h*) is the component size at threshold *h*, and *E* and *H* are the extension and height parameters as defined by Baggio et al (*E*=0.5, *H*=2.25) [37]. Compared to a null TFNBS matrix (creating through group shuffling, n=3,000 permutations), only the observed TFNBS values for p<0.05 are displayed in the end result.

To statistically test differences between networks before and after NTX, an average OUD-only sample was created at pre-NTX and on-NTX (bootstrap 1,000 samples) to have equal N to the co-morbid SUD+OUD group and then pairwise subtraction was performed by pair shuffling (1,000 permutations). Once an average difference matrix was acquired for pre-NTX and on-NTX, the Euclidean norm (ε) [38] was used to quantify differences at each time-point as a singular value per network:

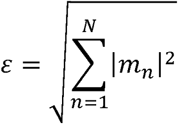

Where *m* is the difference matrix, indexed by value *n* through *N*. Only pairwise ROIs that reached statistical significance via TFNBS were compared in this way, to avoid difference matrices being heavily weighted by unchanged ROI pairs between the compared conditions. A two-sample t-test was then performed on the Euclidean norms at pre-NTX vs. on-NTX.

All non TFNBS corrected two-sample t-tests were corrected for multiple comparisons by dividing the alpha for significance (0.05) by the total number of comparisons being made (i.e., Bonferroni correction) [39].

## 3. Results

### a. At baseline, CanUD+OUD, AUD+OUD, and CocUD+OUD are all associated with unique FC alterations compared to OUD-only. After NTX, these alterations are diminished in CanUD+OUD and AUD+OUD, but not in CocUD+OUD

Following detoxification and prior to administering NTX, N=10 subjects with CanUD+OUD underwent resting state imaging, along with N=25 subjects with OUD-only. By determining the Pearson correlation coefficient between all combinations of ROIs defined by the Power atlas, the FC of inter- and intra-brain networks were calculated (Figure 1A). At roughly 2-weeks post NTX injection, N=10 subjects with CanUD+OUD underwent resting state imaging, along with N=20 controls with OUD-only and the same FC calculation was done on this data (Figure 1B). To statistically compare the two groups at each time-point, a two-sample t-test was performed on each ROI x ROI FC value. To correct for multiple comparisons, a TFNBS approach was used to place more statistical weight on heavily interconnected graph components, with greater differences appreciated between groups pre-NTX (Figure 1C) than on-NTX (Figure 1D).

**Figure 1:**
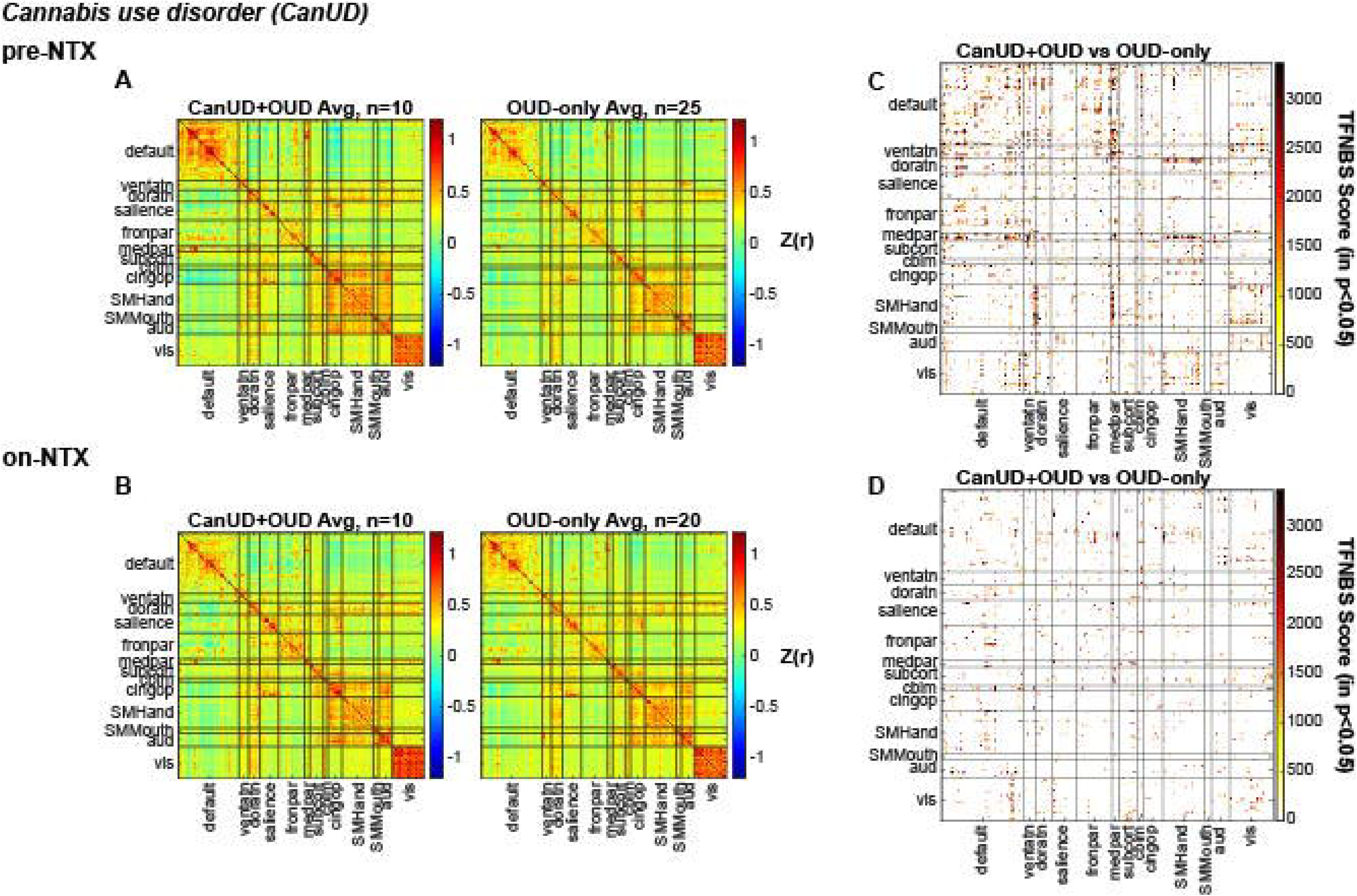
Functional connectivity is altered at baseline in CanUD+OUD compared to OUD-only, but these differences decrease with NTX. Pearson correlation coefficients (r) representing the functional connection strength between two ROI’s within networks specified on the x and y axis at A) baseline and B) after receiving NTX. Matrices are organized to display FC values for (left to right) CanUD+OUD and OUD-only. Matrices displaying the TFNBS scores of CanUD+OUD vs. OUD-only comparisons with p<0.05 by two-sample t-test at C) baseline and D) after receiving NTX.

Using the same experimental timeline and computational model as in Figure 1, the same comparisons were performed using N=6 subjects with AUD+OUD (compared to the same N=25 or N=20 subjects with OUD-only, pre-NTX or on-NTX, respectively, SFigures 2A,B). The same TFNBS approach was used to statistically compare the two groups at each time-point, and similarly to CanUD+OUD in Figure 1, greater differences in FC were found between groups pre-NTX (SFigure 2C) than on-NTX (SFigure 2D). This same analysis was repeated for N=7 subjects with CocUD+OUD (SFigures 3A,B) and did not yield obviously different statistical differences between groups at the two imaging time-points (SFigures 3C,D).

**Figure 2:**
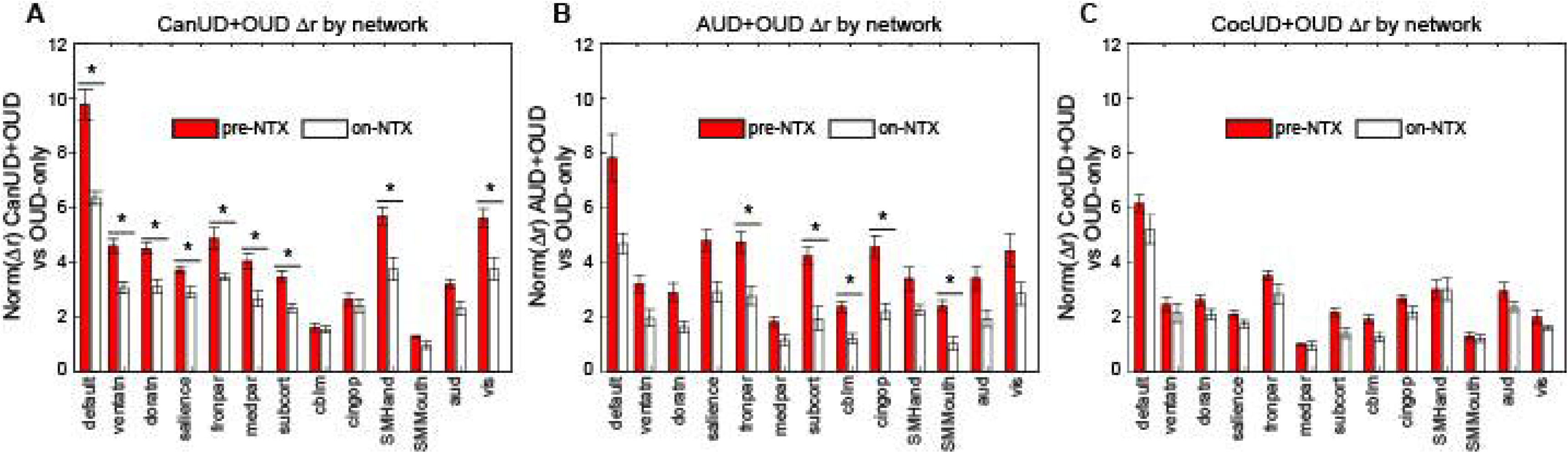
NTX has a normalizing effect on the most networks in CanUD+OUD, followed by AUD+OUD, and hardly any effect in CocUD+OUD. The Euclidean norm of positive differences between A) CanUD+OUD, B) AUD+OUD, and C) CocUD+OUD compared to OUD-only at both time-points. Statistical significance is determined between norm values pre-NTX and on-NTX by two-sample t-test. Significance is determined by alpha=0.05/13 (number of networks).

### b. Network differences compared to OUD-only are significantly reduced with NTX in CanUD+OUD while only changes in parietal, subcortical, sensory, and cerebellar networks are significantly reduced in AUD+OUD. Networks remain relatively unchanged with NTX in CocUD+OUD

There was a striking difference in TFNBS scores between pre-NTX and on-NTX in CanUD+OUD and AUD+OUD compared to OUD-only (Figures 1C,D and SFigures 2C,D), which was better illustrated by taking the network-wise sum of all TFNBS values at each time-point (SFigure 4A-C). For CanUD+OUD and AUD+OUD, the TFNBS scores between pre-NTX and on-NTX decreased by more than half in most networks. This finding was not present in the analysis with CocUD+OUD (SFigure 4C), in fact the total sum of TFNBS remained roughly unchanged between pre-NTX and on-NTX (however there was some re-distribution of which networks demonstrated more/less FC changes compared to OUD-only at the two time-points). Indeed, in CanUD+OUD and AUD+OUD, the difference between these SUD+OUD FC matrices and OUD-only FC matrices pre-NTX were mostly due to hyperconnectivity in the SUD+OUD condition (increased positive correlations in SFigure 5, top row). The differences between CanUD+OUD and AUD+OUD with OUD-only on-NTX were reduced, while multiple differences were still noted in CocUD+OUD (SFigure 5, bottom row). To quantify this, we isolated the positive FC differences (representing hyperconnectivity) between each co-morbid SUD+OUD condition and OUD-only pre-NTX and calculated the Euclidean norm by network (Figure 2, larger norm meaning larger difference between SUD+OUD and OUD-only; smaller norm, smaller difference). Again, the Euclidean norm was taken of the positive differences present on-NTX between each SUD+OUD condition and the OUD-only group. In almost every network the difference between CanUD+OUD and OUD-only decreased on-NTX (Figure 2A). In AUD+OUD, parietal, subcortical, sensory, and cerebellar networks approached the values seen in OUD-only on-NTX (Figure 2B), while no significant change was seen in CocUD+OUD (Figure 2C). While less prominent at baseline, the same analysis was done for negative differences (representing hypoconnectivity) between each SUD+OUD condition and OUD-only, which had more variable results on-NTX (SFigure 6).

### c. In CanUD+OUD, FC alterations within DMN were stronger in younger brains compared to in OUD-only

Notably, the DMN has the highest density of ROIs per the Power atlas, which partially explains the large TFNBS score observed in this network (SFigure 4A). However, in CanUD+OUD, using both the TFNBS and Euclidean norm calculations (the latter not dependent on ROI number), the DMN had the greatest change in FC strength on-NTX (Figure 2A, SFigure 4A). We used this network to illustrate each individual subject trajectory from pre-NTX to on-NTX. The average FC value within DMN was plotted at each time-point and an ANOVA was used to evaluate the interaction of group (CauUD+OUD vs. OUD-only) by time (SFigure 7A). Further, the relationship between average FC in DMN was inversely related to age in the CanUD+OUD group and also in the OUD-only group (SFigures 7B,C). The inverse relationship was stronger in the CanUD+OUD group compared to the OUD only group (R^2^=0.64 vs R^2^=0.12). These results were likely not largely influenced by a differential distribution of age or ASI within groups (SFigures 8A,B).

## 4. Discussion

This study represents secondary analyses to investigate the differential effect of NTX on functional network connectivity in the setting of co-morbid CanUD, AUD, or CocUD. Pearson FC analysis was performed at baseline and roughly two weeks after a therapeutic dose of long-acting injectable NTX. A graph-theory based statistical approach yielded large and significant differences in FC between each SUD+OUD condition and OUD-only pre-NTX that mostly resolved on-NTX in CanUD+OUD and AUD+OUD, but not in CocUD+OUD. While NTX is an established treatment for OUD and AUD, no treatment has been established for CanUD. Specifically, in CanUD+OUD, the default mode network (**DMN**) was the most affected at baseline with bigger FC alterations present in younger subjects.

Functional connectivity is a non-invasive surrogate measure of coordinated neural activity [40], and has been robustly shown to be sensitive to underlying disease processes in humans [28, 30] and animal models [41, 42]. Both degradation and enhancement of FC strength amongst networks can be pathological. In fact, in neuroimaging studies focused on cognitive impairment and unhealthy aging, main findings have dictated a period of functional hyperconnectivity preceding the late-stage hypoconnectivity that is usually found in steep cognitive decline/dementia [43]. In SUDs, recent neuroimaging studies in OUD and AUD have described a degradation of functional network segregation, an expression that quantifies the overall loss of variance in FC values (e.g., globally hyper- or hypo-connected brains) [24, 25]. Fortunately, despite changes in brain architecture, brains have a remarkable ability to rewire and regain function (i.e., plasticity). In otherwise healthy individuals this can occur in as little as 48 hours [29]. While the presented time-course of this study is rather short (∼2 weeks), the inclusion/exclusion criteria resulted in a sample of relatively healthy individuals, outside of their SUD(s), making the two-week time-point a reasonable interval to expect plasticity to occur. In that time, the positive differences (representing hyperconnectivity) between CanUD+OUD and AUD+OUD with OUD-only significantly decreased from pre-NTX to on-NTX (Figures 2A,B) and there were instances of increased negative differences (representing hypoconnectivity) on-NTX (SFigures 6A,B). Pre-NTX, the positive differences outweighed the negative differences, indicating that co-morbid CanUD and AUD induced more global hyperconnectivity than hypoconnectivity when present with OUD, but this hyperconnectivity was responsive to NTX. These measures of hyper- and hypoconnectivity were more uniform in co-morbid CocUD at both time-points (Figure 2C and SFigure 6C). Still, the increase in negative differences (i.e., induced hypoconnectivity) between CanUD+OUD and AUD+OUD with OUD-only on-NTX is notable, and has been noted elsewhere in investigating psilosybin as a treatment for AUD and is hypothesized to be responsible for blunting reward-adjacent pathways [44].

Since the findings related to OUD-only have been previously described [45, 46], the goal of the present analysis was to delineate functional alterations due to co-morbid SUDs, while controlling for opioid use. To this end, all co-morbid SUD comparisons were against OUD subjects without any other co-morbid SUDs, and five subjects were removed based on heavier opioid use compared to the rest of the group (SFigure 1A). Using the statistical methods described, the largest change in FC after NTX was within default mode network in CanUD+OUD. The DMN is foundational in the brain’s functional integrity and internal mentation at rest but also gets activated during episodic memory and predicting future tasks, processes that are frequently disrupted in psychiatric illness (e.g., schizophrenia, depression) [47, 48]. Made up of the ventral medial prefrontal cortex, the dorsal medial prefrontal cortex, and the posterior cingulate cortex, the DMN is spatially disjointed from the dorsal attention network (**DAN**). Also functionally antagonistic, the DAN is primarily indicated in visuospatial attention and therefore utilized in most task-based behaviors. Made up of the intraparietal sulcus and frontal eye fields, the DAN primarily integrates sensory information to coordinate a motor response [49]. A third constellation of brain regions has been shown to be a functional mediator of the DMN and DAN during goal-directed activity [50]. The fronto-parietal network is made up of the dorsolateral prefrontal cortex and posterior parietal cortex and largely thought to be the control center of salience processing and executive function [51]. The fronto-parietal network has been shown to functionally interplay between the DMN and DAN, with nodes aligned with each as well as aligned with both, to complete a triangular unit that modulates an individual’s ability to incorporate and respond to internal and external stimuli [52, 53]. Functional alteration within any of these three networks has been described in patients at high-risk of psychosis [54], experiencing psychotic-like episodes [55], and those without psychosis but with a history of cannabis use [56]. Further, this constellation of networks, as well as interactions with the salience network have been shown to influence drug-taking behaviors and be associated with negative emotions, ruminations, and impaired self-awareness [57]. This is consistent with the findings from behavioral cannabis studies in adolescents showing a decline in working memory, attention, and executive functioning into adulthood [58]. It is concerning that our analysis here shows a larger baseline effect in younger individuals (SFigure 7B), emphasizing the need for earlier intervention for CanUD. The results here are encouraging in that NTX resulted in diminished FC alterations in all aforementioned networks in CanUD+OUD (Figure 2A).

In AUD+OUD, parietal, sensory, and cerebellar were the main networks affected by NTX. Unique to AUD+OUD is the decrease in network differences with OUD-only in cerebellar FC after NTX. In AUD, multiple studies document cerebellar dysfunction both in acute intoxication from alcohol and in chronic AUD. From in-utero development, where exposure to alcohol induces multiple cerebellar deficits [59], to the acute intoxication period in adolescence/adulthood, where high blood alcohol levels start to impact functions governed by the cerebellum (e.g., coordination), to long term use in AUD, ethanol impacts GABA signaling in Purkinje cells as well as interneurons and granule cells that fortify the neural transmission pathways of the cerebellum [60]. Elsewhere, connectivity with cerebellar networks predicted response to brief interventions with NTX in AUD [61], which agrees with the result presented here. The baseline FC alterations in CocUD+OUD compared to OUD-only were less pronounced than in CanUD+OUD and AUD+OUD, but still present (SFigure 3C). In a recent study, CocUD manifested in a unique FC signature that was distinguishable from controls, involving inter-network connections in frontoparietal, default mode, dorsal attention, limbic, ventral attention, visual, and somatomotor networks, which was also evidenced here (SFigures 3C and 4C) [62]. Unfortunately, treatment with NTX did not mitigate any of these findings. Although there were less negative differences present within visual network between CocUD+OUD and OUD-only on-NTX (SFigure 6C), the overall TFNBS score was greater on-NTX than pre-NTX for visual network (SFigure 4C), indicating that while more ROIs became similar between CocUD+OUD and OUD-only on-NTX (smaller Euclidean norm), there were still individual ROIs that were significantly altered that were within largely inter-connected networks (greater TFNBS score).

As mentioned above, there is no FDA approved treatment of CanUD. NTX, however, is approved for the treatment of both OUD and AUD. The mechanism by which NTX addresses alcohol misuse is through decreasing the perceived reward associated with drinking [17]. This decreased reward has also been reported in CanUD with NTX [21], though the effectiveness of NTX for CanUD needs further investigation. Notably, NTX has been ineffective for CocUD, [18]. Therefore, the present imaging results in AUD+OUD and CocUD+OUD are consistent with the current clinical evidence for NTX to treat either disorder; with AUD, NTX mitigated functional network alterations, in CocUD, it seemed to have little effect. The results in CanUD+OUD were promising, as FC alterations were significantly mitigated after a single dose, similar to the response seen in AUD+OUD. While the ideal treatment for a SUD would involve long-term maintenance of network activity indicated in addiction (e.g., reward, executive, and memory networks), the ability for NTX to affect these networks long-term is outside the scope of this work. Instead, we present further evidence that NTX can aide to acutely restore some of the early functional changes present in patients with OUD and co-morbid CanUD.

## 5. Conclusions

Here, we performed FC analysis on subjects with OUD and co-morbid CanUD, AUD, or CocUD. We analyzed networks before and after receiving a long-acting form of NTX and note significant changes in functional architecture induced by NTX for subjects with co-morbid CanUD and AUD, but not CocUD. For subjects with co-morbid CanUD, younger subjects had greater baseline effects within the DMN, indicating a potential need for earlier intervention in this demographic. Finally, we provide further evidence of NTX being a potentially efficacious treatment for CanUD.

## Acknowledgements

This work was supported by the Commonwealth of Pennsylvania CURE Addiction Center of Excellence grant (#4100055577, DDL), and the following National Institutes of Health grants: R25MH119043 (LMB), R01DA036028 (DDL), AA031088 (CEW), AA031337 (CEW), AA031570 (CEW), and K01DA051709 (ZS).

---

**SFigure 1:**
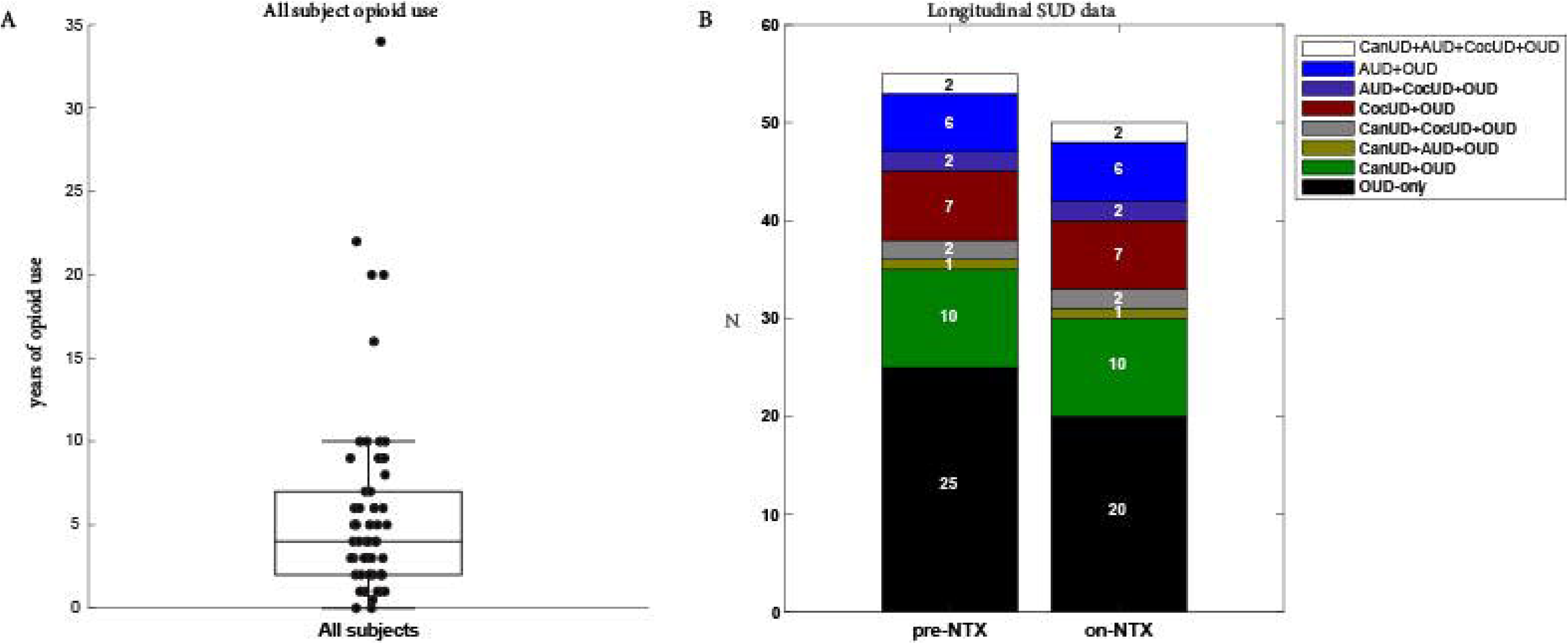
SUDs within imaged subjects. A) Years of opioid use in all subjects considered for analysis. The 5 statistical outliers were removed from the analysis reported in this study. B) Number of subjects with each substance use disorder (or combination of disorders) at each time-point.

**SFigure 2:**
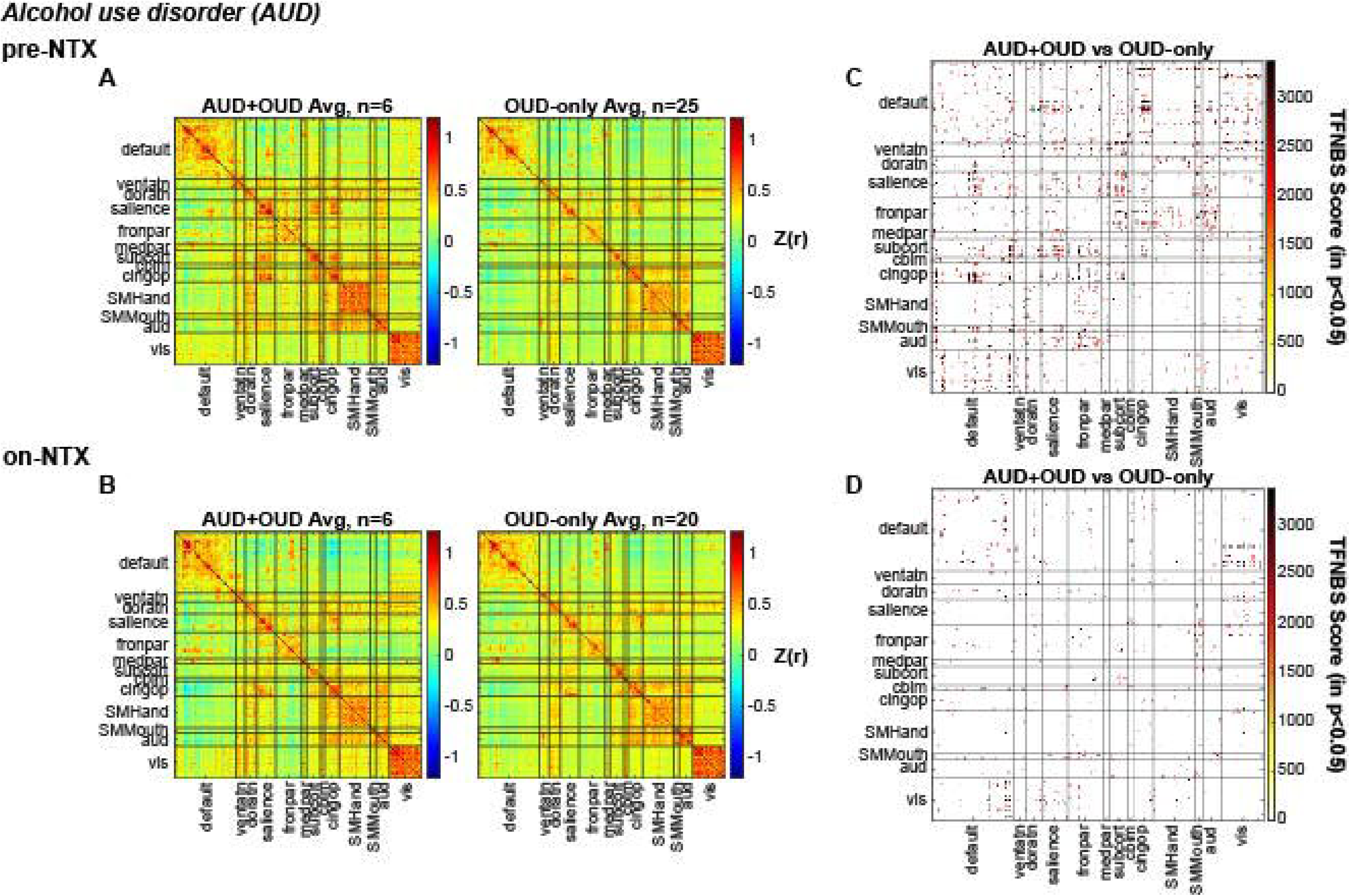
Functional connectivity is altered at baseline in AUD+OUD compared to OUD-only, but these differences decrease with NTX, as expected. Pearson correlation coefficients (r) representing the functional connection strength between two ROI’s within networks specified on the x and y axis at A) baseline and B) after receiving NTX. Matrices are organized to display FC values for (left to right) AUD+OUD and OUD-only. Matrices displaying the TFNBS scores of AUD+OUD vs. OUD-only comparisons with p<0.05 by two-sample t-test at C) baseline and D) after receiving NTX.

**SFigure 3:**
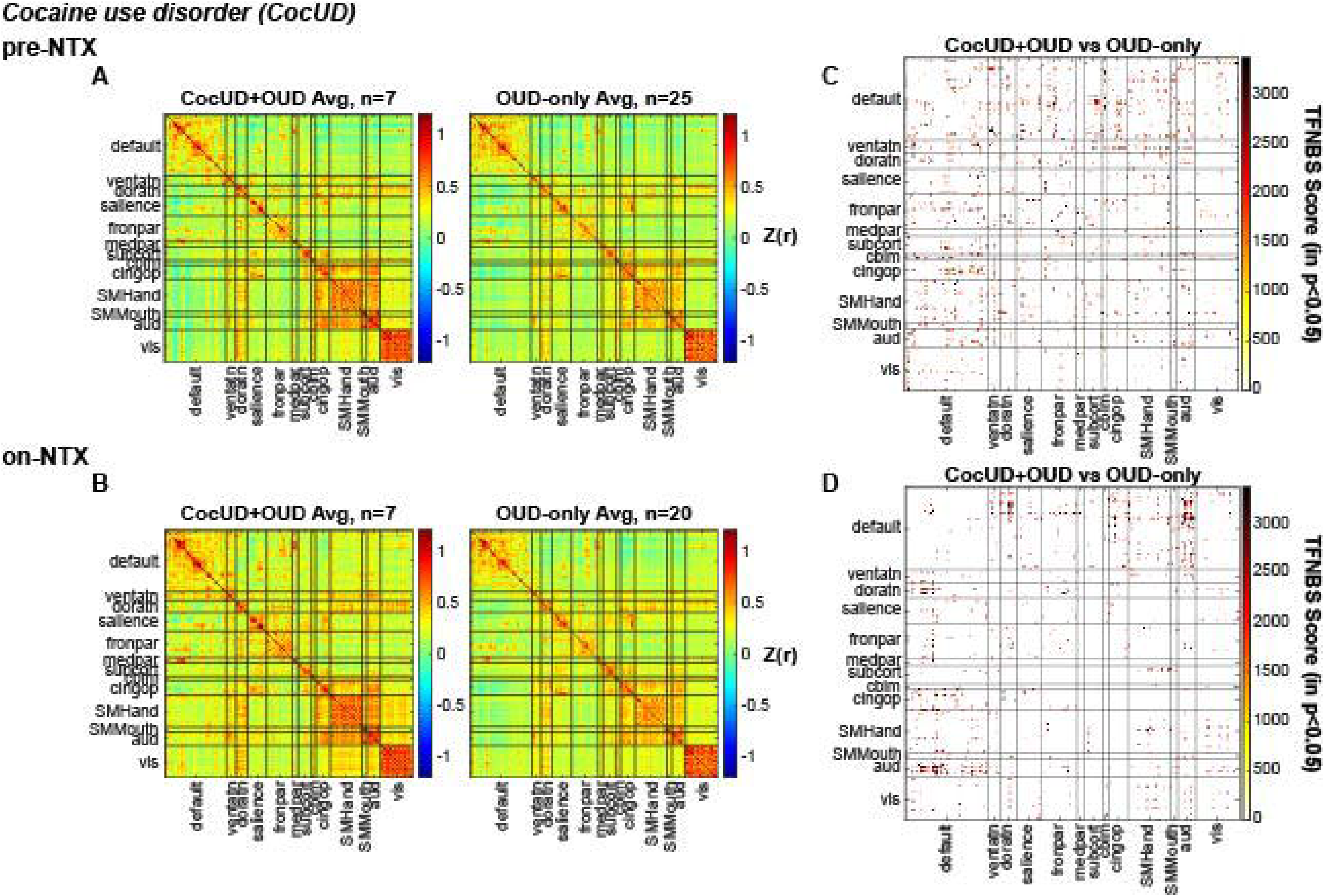
Functional connectivity is altered at baseline in CocUD+OUD compared to OUD-only and is minimally affected by NTX. Pearson correlation coefficients (r) representing the functional connection strength between two ROI’s within networks specified on the x and y axis at A) baseline and B) after receiving NTX. Matrices are organized to display FC values for (left to right) CocUD+OUD and OUD-only. Matrices displaying the TFNBS scores of CocUD+OUD vs. OUD-only comparisons with p<0.05 by two-sample t-test at C) baseline and D) after receiving NTX.

**SFigure 4:**
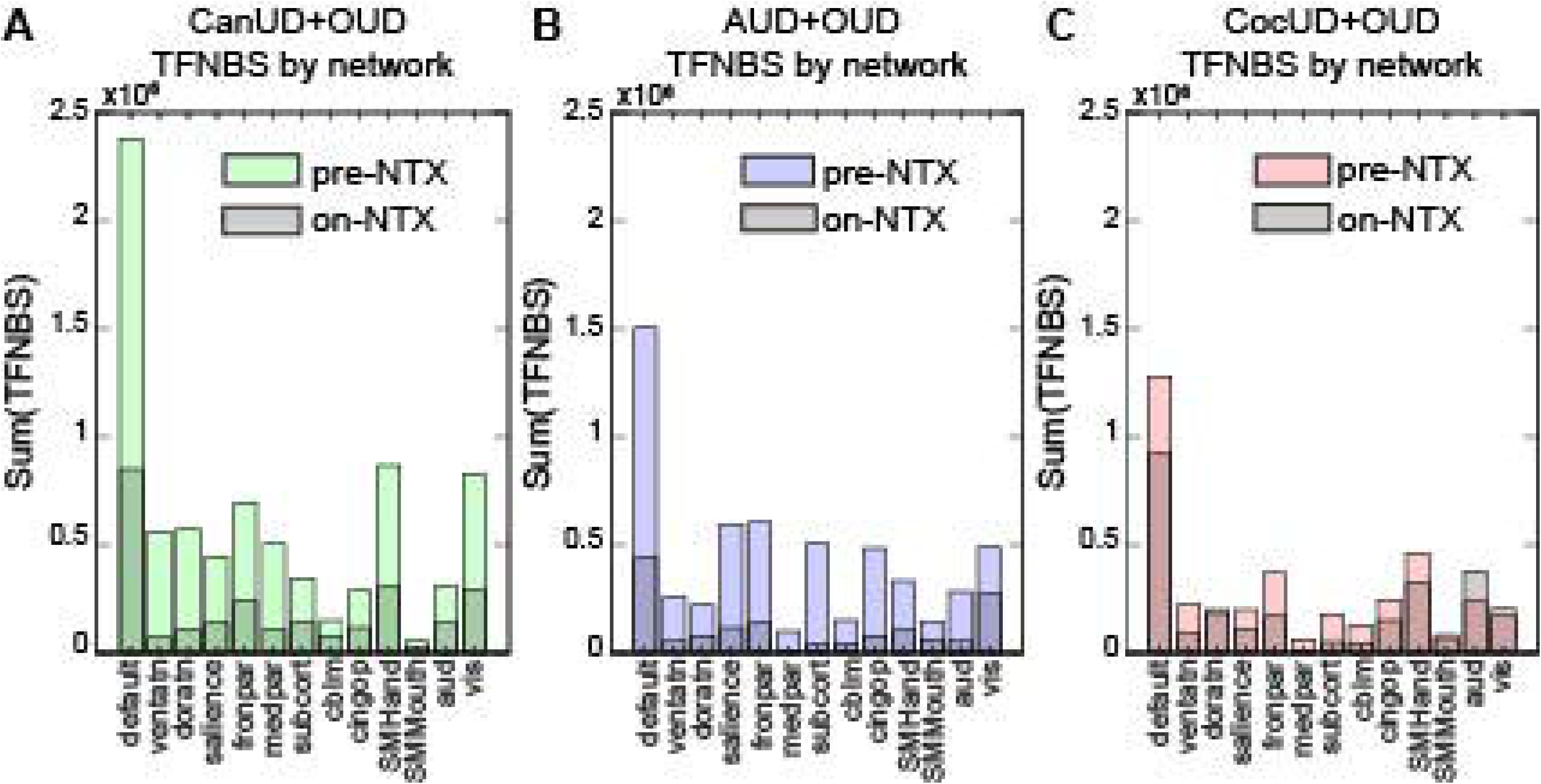
FC differences between CanUD+OUD and AUD+OUD with OUD-only decrease with NTX, but not in CocUD+OUD. The within-network sum of TFNBS scores that result from comparing each co-morbid SUD+OUD vs OUD-only at baseline (in color) or after NTX (black) in A) CanUD+OUD B) AUD+OUD and C) CocUD+OUD.

**SFigure 5:**
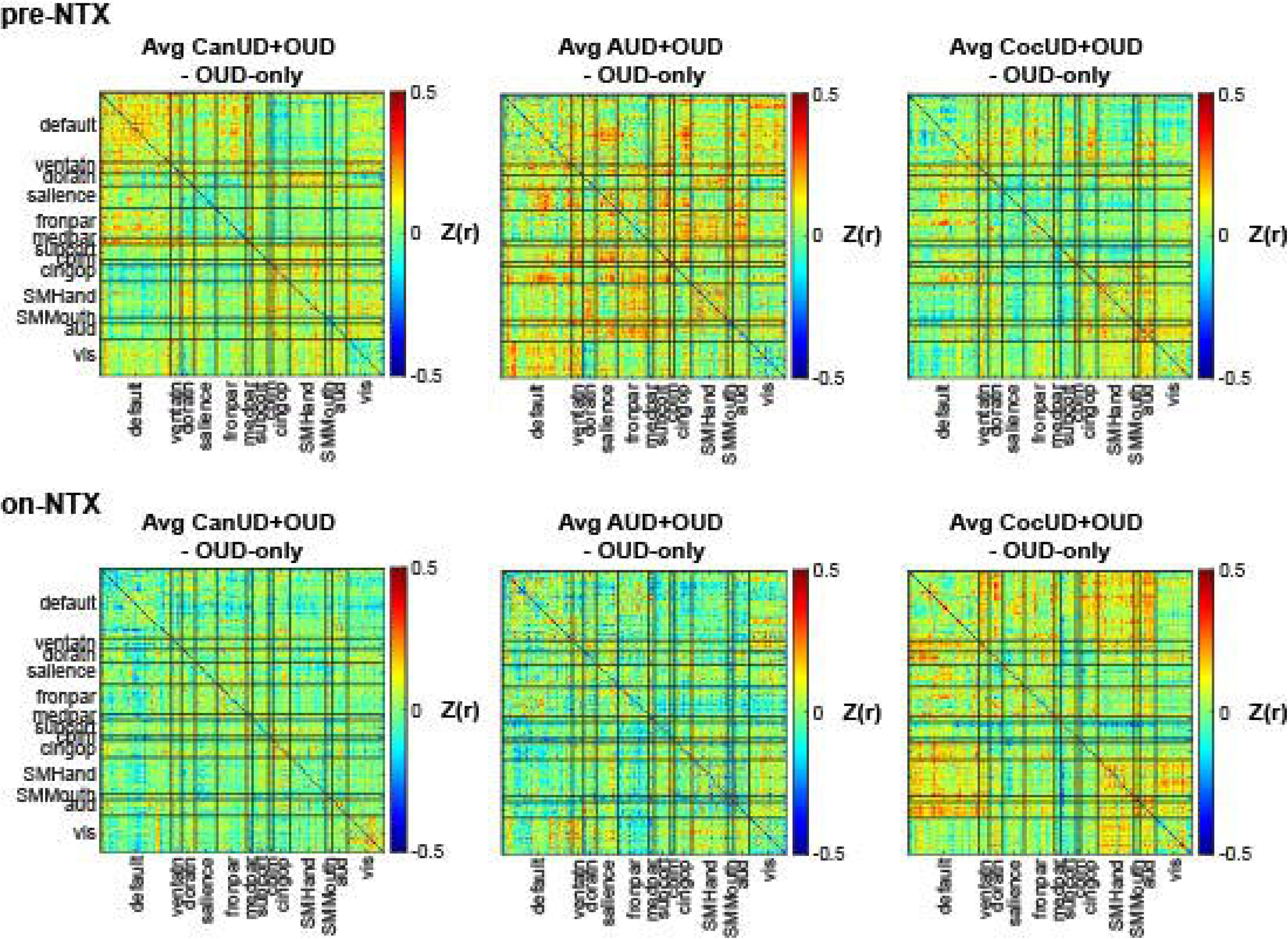
Difference matrices between each SUD+OUD condition and OUD-only either pre-NTX or on-NTX.

**SFigure 6:**
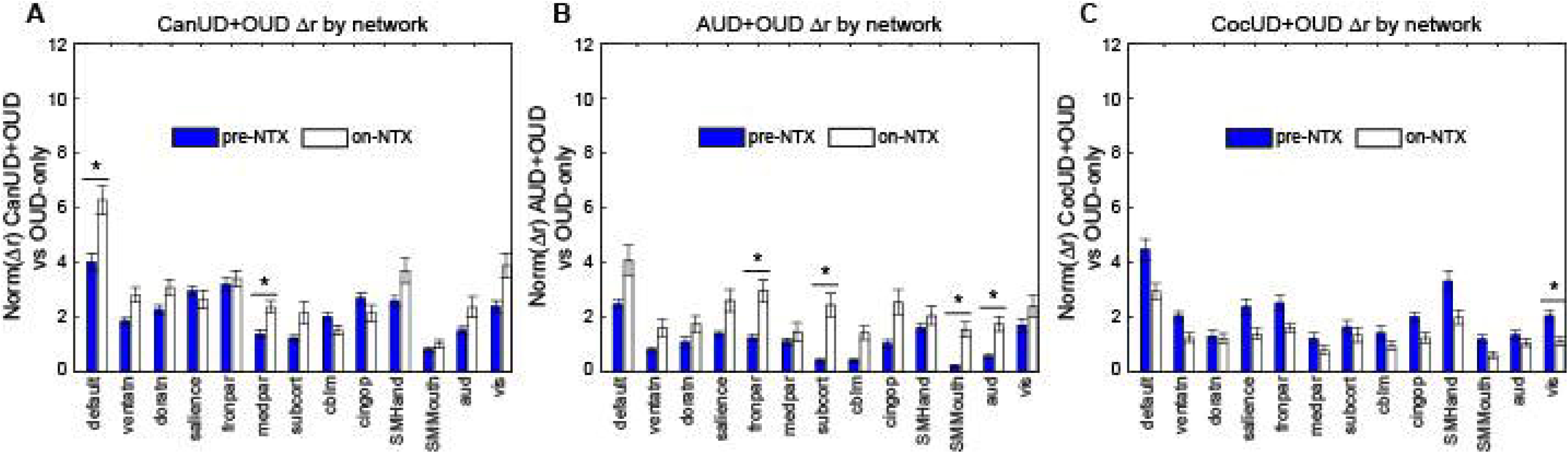
NTX results in hypoconnectivity in isolated networks in CanUD+OUD and AUD+OUD, but not CocUD+OUD. The Euclidean norm of negative differences between A) CanUD+OUD, B) AUD+OUD, and C) CocUD+OUD compared to OUD-only at both time-points. Statistical significance is determined between norm values pre-NTX and on-NTX by two-sample t-test. Significance is determined by alpha=0.05/13 (number of networks).

**SFigure 7:**
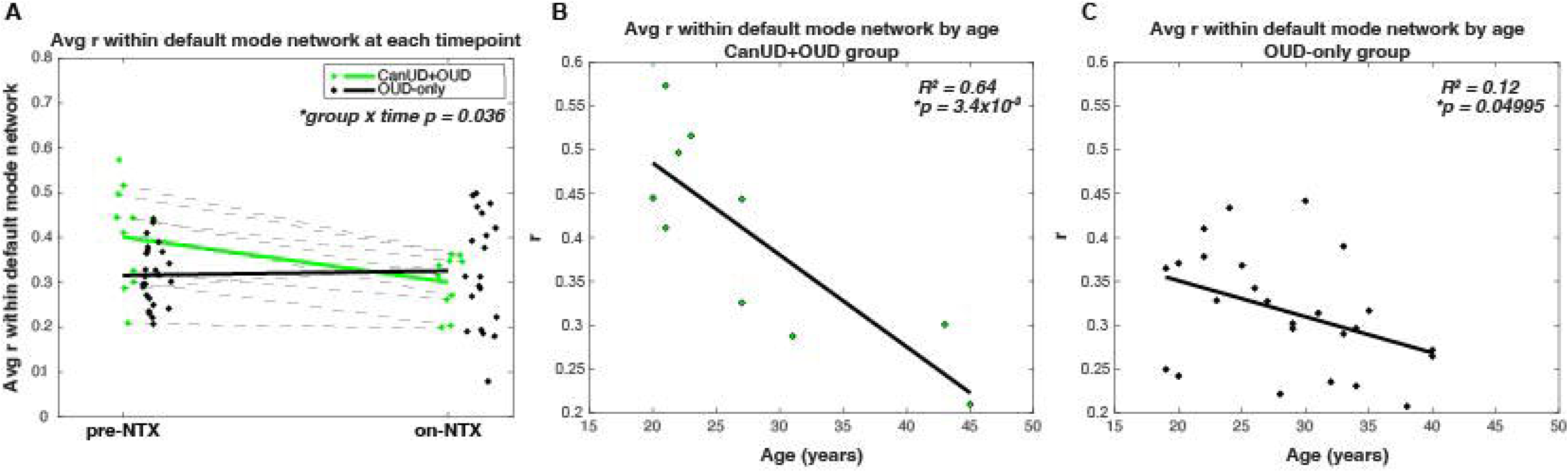
In default mode network, younger subjects had bigger FC alterations at baseline in those with CanUD+OUD. A) Average Pearson correlation coefficient (r) within default mode network for each subject (dots) with random horizontal scatter for visibility. Dotted lines connect subjects at each time-point and solid lines represent the group average at each time-point. Statistical comparison done by ANOVA with significant interaction between group and time-point. Linear regression between average Pearson correlation coefficient (r) within default mode network and subject age in B) CanUD+OUD and C) OUD-only.

**SFigure 8:**
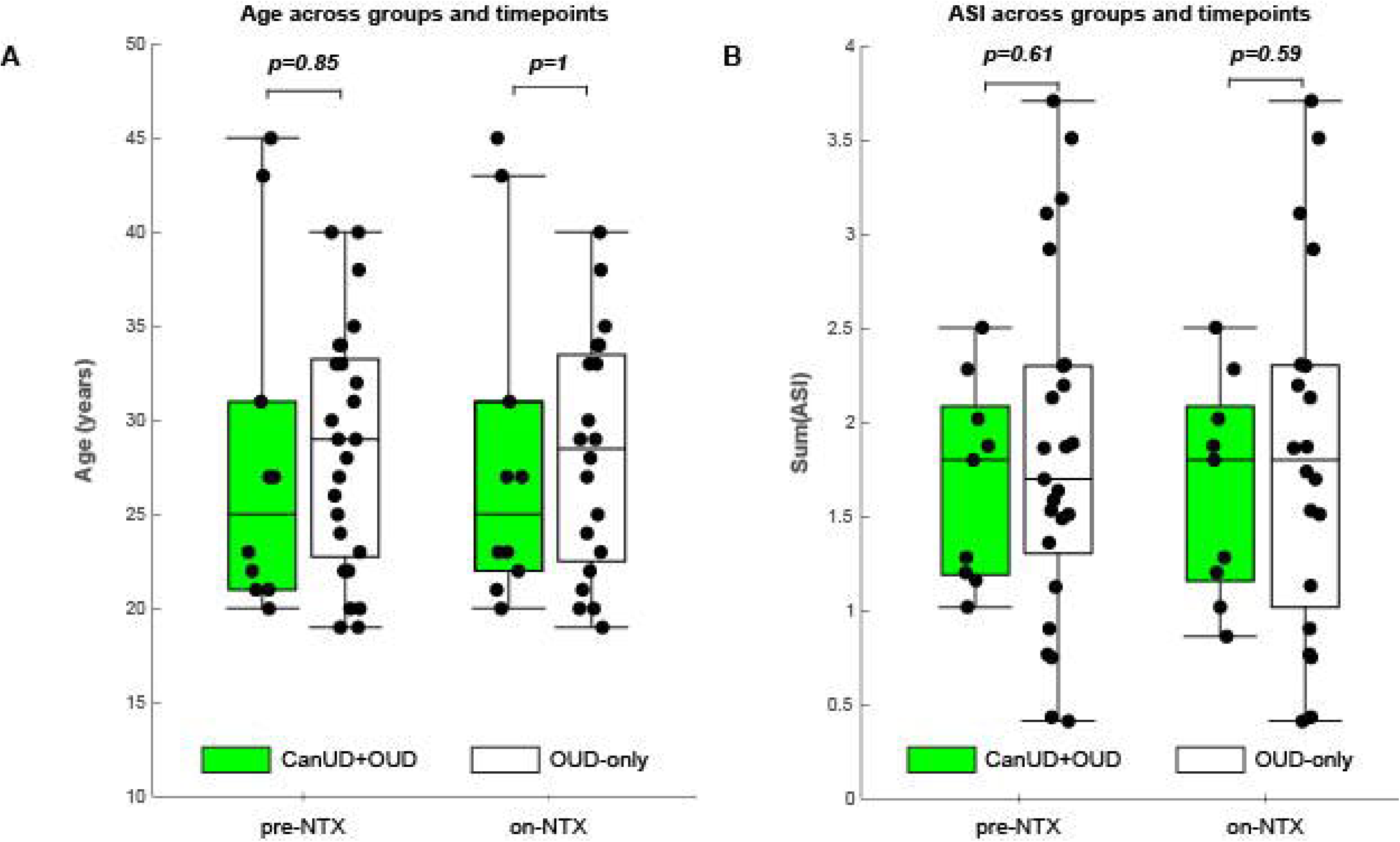
Age and ASI severity is evenly distributed between CanUD+OUD and OUD-only groups at each time-point. Each subject’s A) age or B) summed ASI score (dots) at each time-point with boxplots overlaid to represent median, interquartile range, and minimum/maximum values not considered outliers. P-values represent two-sample t-tests between groups at each time-point (NS).

